# RNAmining: A machine learning stand-alone and web server tool for RNA coding potential prediction

**DOI:** 10.1101/2020.10.26.354357

**Authors:** Thaís A. R. Ramos, Nilbson R. O. Galindo, Raúl Arias-Carrasco, Cecília F. da Silva, Vinicius Maracaja-Coutinho, Thaís G. do Rêgo

## Abstract

Non-coding RNAs (ncRNAs) are important players in the cellular regulation of organisms from different kingdoms. One of the key steps in ncRNAs research is the ability to distinguish coding/non-coding sequences. We applied 7 machine learning algorithms (Naive Bayes, SVM, KNN, Random Forest, XGBoost, ANN and DL) through 15 model organisms from different evolutionary branches. Then, we created a stand-alone and web server tool (RNAmining) to distinguish coding and noncoding sequences, selecting the algorithm with the best performance (XGBoost). Firstly, we used coding/non-coding sequences downloaded from Ensembl (April 14th, 2020). Then, coding/non-coding sequences were balanced, had their tri-nucleotides counts analysed and we performed a normalization by the sequence length. Thus, in total we built 180 models. All the machine learning algorithms tests were performed using 10-folds cross-validation and we selected the algorithm with the best results (XGBoost) to implement at RNAmining. Best F1-scores ranged from 97.56% to 99.57% depending on the organism. Moreover, we produced a benchmarking with other tools already in literature (CPAT, CPC2, RNAcon and Transdecoder) and our results outperformed them, opening opportunities for the development of RNAmining, which is freely available at https://rnamining.integrativebioinformatics.me/.

## INTRODUCTION

Non-coding RNAs (ncRNAs) are key functional players on different biological processes in organisms from all domains of life (1,2). Its investigation is already routine in almost every transcriptome or genome project. Dysregulations in these molecules may lead to different types of human disease, including cancers (3), neurological disorders (4) and cardiovascular infirmities (5).

The genome of eukaryotic (6) organisms is, in general, composed majoritary by non-coding transcripts, with complex organisms estimated to transcribe more than 75% of their genomes (7). Besides strong evidence associating these ncRNAs to key functions in the cell, their majority are not yet associated with a functional mechanism. In a transcriptome project exists an important step in the computational identification of ncRNAs, which is the evaluation of their potential to be translated into proteins using different bioinformatics approaches (8,9). To computationally evaluate the coding potential of a set of transcripts, available tools or algorithms normally analyse specific characteristics available in its primary sequences (*e.g.* nucleotides counts, the existence of a trustful open reading frame).

For instance, RNAcon implements a SVM-based model for the discrimination between coding and non-coding sequences (10). Coding Potential Assessment Tool (CPAT) (11) assesses the coding potential through an alignment-free method, which uses a logistic regression model built based on different characteristics of the sequence open reading frame (ORF), which includes length, coverage and nucleotides compositional bias. TransDecoder identifies candidate coding transcripts based on other distinctive features from predicted ORFs (*e.g*. a minimum length ORF, a log-likelihood score, encapsulated ORF) (12). CPC2 (13) trained a support vector machine (SVM) model using Fickett TESTCODE score, open reading frame (ORF) length, ORF integrity and isoelectric point as features. The LIBSVM (14) package was employed by training a SVM model using the standard radial basis function kernel (RBF kernel) with the training dataset containing 17,984 high-confident human proteincoding transcripts and 10,452 non-coding transcripts (11).

Here, we applied and benchmarked seven different machine learning algorithms (Random Forest, Gradient boosting (XGBoost), Naive Bayes, K-Nearest Neighbors (K-NN), Support Vector Machine (SVM), Artificial Neural Network (ANN) and Deep Learning (DL)) through 15 organisms from different evolutionary branches, in order to evaluate their performance in distinguishing coding and non-coding RNA sequences. Next, we developed a stand-alone and web server tool, called RNAmining (http://rnamining.integrativebioinformatics.me/), by selecting and implementing the algorithm with the best performance in all organisms (XGBoost). RNAmining was evaluated through 24 organisms from the eukaryotic tree of life and its results outperformed public available tools commonly used for that purpose.

## MATERIAL AND METHODS

### Machine learning classifier algorithms selection

In the classification process there is a division related to the learning paradigm, with classification algorithms divided into: (i) *Symbolic,* which seeks to learn by constructing symbolic representations of a concept through the analysis of examples and counterexamples (e.g. Decision Trees and Rule-based System); (ii) *Statistical,* which looks for statistical methods and use models to find a good approximation of the induced concept (*e.g.* Bayesian learning); (iii) *Based on Examples* (lazy systems), which aims to classify examples never seen using similar known examples, assuming that the new example will belong to the same class as the similar example (*e.g*. K-Nearest neighbor); (iv) *Based on Optimization,* which consists of maximizing (or minimizing) an objective function or finding an optimal hyperplane that best divides two classes (*e.g.* Support Vector Machine (SVM) and Neural Networks); (v) *Connectionist Representation,* which represents simplified mathematical constructions inspired by the biological model of the nervous system (e.g. Neural Networks). In this benchmarking, we decided to evaluate the performance of selected algorithms from each paradigm type in the coding potential prediction of RNA sequences: Random Forest, XGBoost, Naive Bayes, K-NN, SVM and Neural Networks (Artificial Neural Networks and Convolutional Neural Networks (CNN)).

All the machine learning methods were executed using scikit-learn (Version 0.21.3) (15), except for Neural Network and Deep Learning models which were implemented using Keras API with Tensorflow as backend (Version 2.3.0) (https://github.com/keras-team/keras) and XGBoost algorithm which was executed using XGBoost Library (version 1.2.0) (16) in Python Language (Version 3.8). The Random Forest model was implemented using empirical tests and the best result was selected for training the model. We considered the default parameters with the exception of the number of trees used (150 estimators) and the criterion parameter setted to ‘entropy’ for information gain. KNN and Naive Bayes models were trained with the default values. The SVM parameters were obtained through grid search method and the resulting model was trained with the Radial Basis Function (RBF) kernel, with the Regularization parameter (C) and Kernel coefficient (Gamma) defined in 1000 and 0.8, respectively. Artificial Neural Networks and Deep Learning were performed with different architectures according to grid search and empirical tests. The first ANN experiment was composed of three hidden layers consisting of 32-16-8 neurons, respectively; the second ANN experiment was performed with 64-32-16-8 neurons; and the third experiment was executed with 32-32-16-8 neurons. Next, we produced four experiments with Deep Learning using 2 CNN layers, followed by 2 fully connected (dense) layers: the first experiment had 512(CNN)-512(CNN) filters and 28(Dense)-1(Dense) neurons; the second was created with 64(CNN)-64(CNN) filters and 128(Dense)-1(Dense) neurons; the third was performed with 32(CNN)-32(CNN)-128(Dense)-1(Dense) neurons; and the last was built with 128(CNN)-128(CNN)-128(Dense)-1(Dense) neurons. These layers received as input the total number of attributes (i.e. combination of tri-nucleotides counts, described in the next topics). The hyperparameters used to execute the DL and ANN approaches are made available in Supplementary File S1.

### Datasets selection and filtering criteria

We compared the algorithms performances using different sets of coding and non-coding RNA sequences from Ensembl (April 14th 2020) (17) database, covering 15 organisms of distinct representative Chordata clades (Figure 1A): *Anolis carolinensis* (Sauria, Squamata)*, Chrysemys picta bellii* (Sauria, Testudines)*, Crocodylus porosus* (Archosauria, Pseudosuchia)*, Danio rerio* (Actinopterygii, Teleostei)*, Eptatretus burgeri* (Agnatha, Myxinidae)*, Gallus gallus* (Archosauria, Theropoda)*, Homo sapiens* (Placentalia)*, Latimeria chalumnae* (Sarcopterygii, Coelacanth)*, Monodelphis domestica* (Marsupialia)*, Mus musculus* (Placentalia)*, Notechis scutatus* (Sauria, Squamata)*, Ornithorhynchus anatinus* (Monotremata)*, Petromyzon marinus* (Agnatha, Petromyzontiformes)*, Sphenodon punctatus* (Sauria, Rhynchocephalia)*, Xenopus tropicalis* (Amphibia). All non-coding RNA sequences for each organism were downloaded from Ensembl transcripts. In order to obtain a balanced set of sequences (*i.e.* equal number of coding and noncoding), the group of coding RNAs were randomly selected in order to obtain the same number of ncRNAs for each species. Moreover, before generating the models, the sequences were normalized through their length (*i.e.* each tri-nucleotide count was divided by the total size of the given sequence). All sequences in FASTA format can be retrieved at RNAmining website (https://rnamining.integrativebioinformatics.me/download).

**Figure 1.**
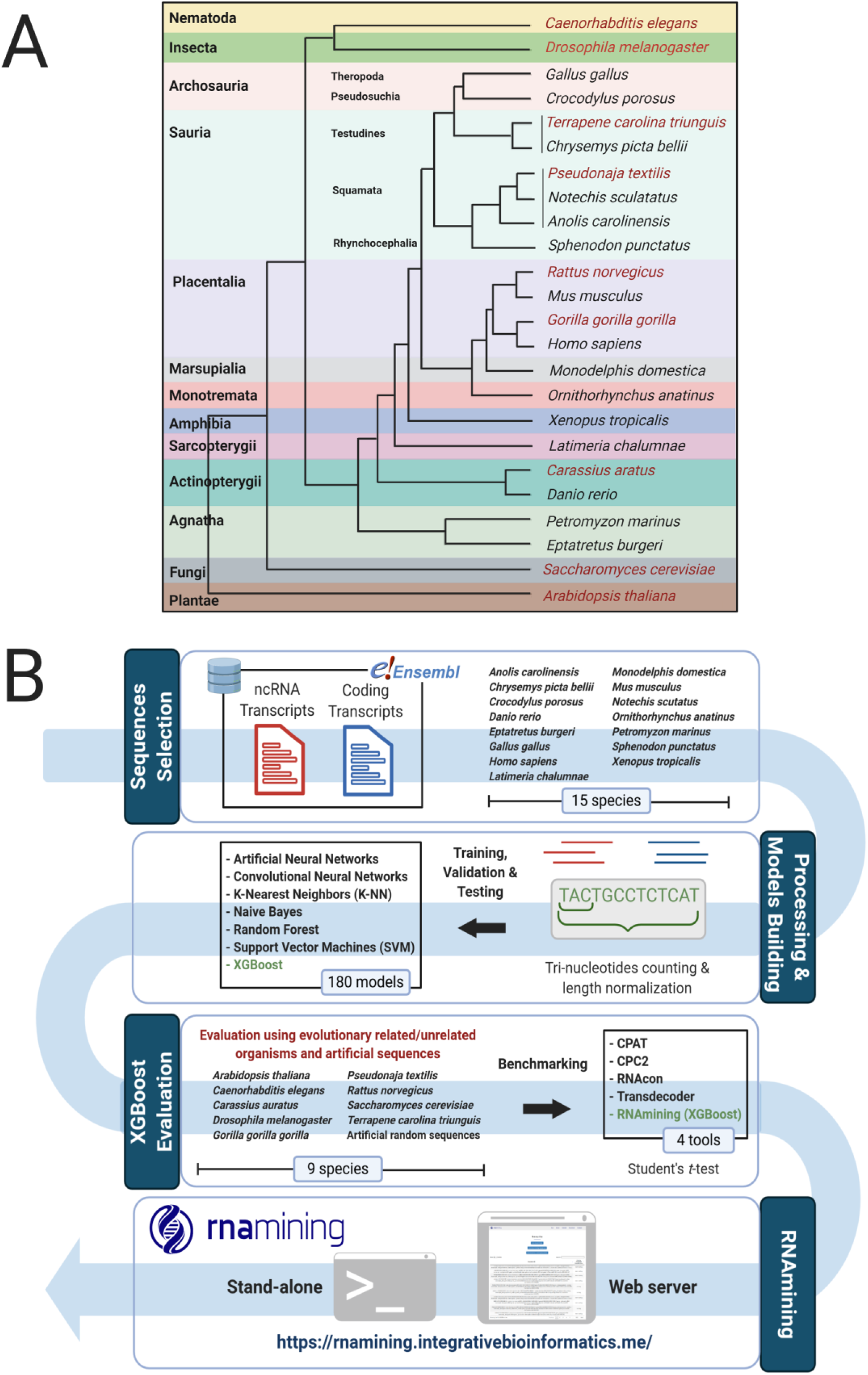
**A. Taxonomic tree according to the used organisms for the models building (black color) and validation (red color). B. Pipeline used to perform the benchmarking and create the tool.** Firstly, we download the coding and non-coding sequences from Ensembl; Next, we perform the trinucleotides counts and sequence normalization. After this, we created a machine learning benchmarking within the 7 algorithms and selected the one with the best performance to be implemented in the RNAmining tool (XGBoost algorithm), which was again evaluated using sequences from 9 other different species and sets of artificially generated ones. Finally, we performed a novel benchmarking with RNAmining against the public available tools for coding potential prediction.

### Training and testing datasets, model building and quality measuring for coding potential evaluation

Sequences were randomly divided into training and testing datasets, using 80% of the data for training and 20% for testing. For ANN and CNN experiments, sequences were splitted into 60% of the data for training and 20% for validation. The testing dataset was the same used in the other machine learning algorithms. The number of sequences used for each organism for the training and test sets can be observed in Table 1. Next, we generated 180 models (*i.e.* one per algorithm for each organism, whereas three experiments for ANN models and four experiments for CNN models), which were further evaluated in this work.

After selection of the best model, it was applied and evaluated in other nine organisms (Figure 1A), including five Chordata and four phylogenetically distant species. Among the chordates, the models were tested in *Carassius auratus* (Actinopterygii, Teleostei), *Gorilla gorilla gorilla* (Placentalia), *Pseudonaja textilis* (Sauria, Squamata)*, Rattus norvegicus (*Placentalia,) and *Terrapene carolina triunguis* (Sauria, Testudines). Within non-chordates species, we evaluated the model in *Arabidopsis thaliana* (Plantae, Eudicots), *Caenorhabditis elegans* (Nematoda), *Drosophila melanogaster* (Insecta, Diptera) and *Saccharomyces cerevisiae* (Fungi, Ascomycota). Finally, it was evaluated using artificial sequences containing the same nucleotides composition of the ncRNAs for each species of the testing dataset (Table 1). Ten sets of random sequences containing the same number of ncRNAs per species were generated using MEME suite Version 5.1.1 with default parameters (18). All sequences in FASTA format can be retrieved at RNAmining website (https://rnamining.integrativebioinformatics.me/download).

**Table 1.** Set of sequences used in the training and testing datasets. List of organisms and the total number of sequences used for testing and training both coding and non-coding RNAs. The numbers are separated into training / testing values. All sequences can be retrieved at RNAmining website (https://rnamining.integrativebioinformatics.me/download).

| Species | Total | Coding | ncRNAs |
| --- | --- | --- | --- |
| **Models Generation (training / testing):** |  |  |  |
| *Anolis carolinensis* | 12,542 / 3,136 | 6,243 / 1,596 | 6,299 / 1,540 |
| *Chrysemys picta bellii* | 11,260 / 2,816 | 5,626 / 1,412 | 5,634 / 1,404 |
| *Crocodylus porosus* | 7,388 / 1,848 | 3,700 / 918 | 3,688 / 930 |
| *Danio rerio* | 12,984 / 3,246 | 6,527 / 1,588 | 6,457 / 1,658 |
| *Eptatretus burgeri* | 1,742 / 436 | 867 / 222 | 875 / 214 |
| *Gallus gallus* | 16,851 / 4,213 | 8,426 / 2,106 | 8,425 / 2107 |
| *Homo sapiens* | 92,844 / 23,212 | 46,575 / 11,453 | 46,269 / 11,759 |
| *Latimeria chalumnae* | 4,668 / 1,168 | 2,344 / 574 | 2,324 / 594 |
| *Monodelphis domestica* | 34,336 / 8,584 | 17,113 / 4,347 | 17,223 / 4,237 |
| *Mus musculus* | 35,272 / 8,818 | 17,668 / 4,377 | 17,604 / 4,441 |
| *Notechis scutatus* | 2,705 / 677 | 1,351 / 340 | 1,354 / 337 |
| *Ornithorhynchus anatinus* | 12,604 / 3,152 | 6,280 / 1,598 | 6,324 / 1,554 |
| *Petromyzon marinus* | 4,243 / 1,061 | 2,107 / 545 | 2,136 / 516 |
| *Sphenodon punctatus* | 1,456 / 364 | 723 / 187 | 733 / 177 |
| *Xenopus tropicalis* | 2,224 / 556 | 1,120 / 270 | 1,104 / 286 |
| **RNAmMining Evaluation:** |  |  |  |
| *Arabidopsis thaliana* | 11,308 | 5,654 | 5,654 |
| *Caenorhabditis elegans* | 50,558 | 25,279 | 25,279 |
| *Carassius auratus* | 15,004 | 7,502 | 7,502 |
| *Drosophila melanogaster* | 31,808 | 15,904 | 15,904 |
| *Gorilla gorilla gorilla* | 15,978 | 7,989 | 7,989 |
| *Pseudonaja textilis* | 1,486 | 743 | 743 |
| *Rattus norvegicus* | 18,662 | 9,331 | 9,331 |
| *Saccharomyces cerevisiae* | 848 | 424 | 424 |
| *Terrapene carolina triunguis* | 2,054 | 1,027 | 1,027 |

### Comparisons with publicly available tools

The performance of all algorithms in the coding potential evaluation was compared with publicly available tools commonly employed for this purpose (RNAcon (10), CPAT (11), Transdecoder (12) and CPC2 (13)), using default parameters. It is worth to note that CPAT only made available models for *H. sapiens* with a coding probability (CP) cutoff of 0.364 (*i.e.* CP >=0.364 indicates coding sequence); *M. musculus* with a CP cutoff of 0.44; *D. melanogaster* with a CP cutoff of 0.39; and *D. rerio* with CP cutoff 0.38. Therefore, for the other organisms we built new models using our training sets and we used the statistical method provided by the authors to calculate the cutoffs probability for coding prediction: *A. carolinensis* (0.4); *C. picta bellii* (0.57); *C. porosus* (0.38); *E. burgeri* (0.35); *G. gallus* (0.42); *L. chalumnae* (0.365); *M. domestica* (0.51); *N. scutatus* (0.15); *O. anatinus* (0.28); *P. marinus* (0.34); *S. punctatus* (0.18); *X. tropicalis* (0.25). The whole workflow of RNAmining development can be visualized in Figure 1B.

### Comparisons with publicly available tools

The XGBoost method was implemented using XGBoost Library (version 1.2.0) in Python Language (Version 3.8) and the models for each species were saved using pickle Python’s library. The web server interface was developed using HTML and CSS. The connection within the front and back-end was implemented through Javascript. The control of files and the connection with Python’s scripts was performed through PHP language. RNAmining user friendly tool and its stand-alone version can be accessed at https://rnamining.integrativebioinformatics.me/. Instructions on how to use it and a whole documentation are made available. Its source code with a Docker platform can be freely obtained at https://gitlab.com/integrativebioinformatics/RNAmining.

## RESULTS

### Using machine learning algorithms to improve the coding potential prediction of RNA sequences

It is known that the algorithms performance in predictive analysis is influenced by particularities available in the genomes sequences of the organisms used in the training set (19), and it should be taken into account when developing novel tools for nucleotides coding prediction. Thus, it is necessary to test several methods to observe which ones can have a good prediction for specific species from evolutionary branches. Similar to Panwar *et al*. (10), we used the tri-nucleotides counts to distinguish coding and non-coding sequences. We evaluated the performance of seven machine learning algorithms using representative organisms from different different branches of the Chordata clade. For that, we used a training and testing set composed by sequences from the same species. The algorithm with best performance within all evaluated organisms, according to F1-scores metric, was XGBoost, as one can see in the following: *A. carolinensis* (98.79); *C. picta bellii* (98.00); *C. porosus* (98.15); *D. rerio* (97.98); *E. burgeri* (97.56); *G. gallus* (99.24); *H. sapiens* (98.50); *L. chalumnae* (99.57); *M. domestica* (98.84); *M. musculus* (97.73); *N. scutatus* (96.51); *O. anatinus* (97.61); *P. marinus* (99.42); *S. punctatus* (99.20); *X. tropicalis* (99.13) (Table 2). As observed, XGBoost algorithm presented F-score values above 97.00, with the worst performance obtained for *Eptatretus burgeri* with a F-score of 97.56. The best performance was obtained for *Petromyzon marinus* with 99.42. All detailed performances with sensitivity, specificity, precision, accuracy, F1-score and the confusion matrix from each algorithm is listed in Supplementary File S2. Based on these results, XGBoost was selected to be implemented in a novel web server and stand-alone tool for RNA coding potential prediction called RNAmining.

**Table 2.**
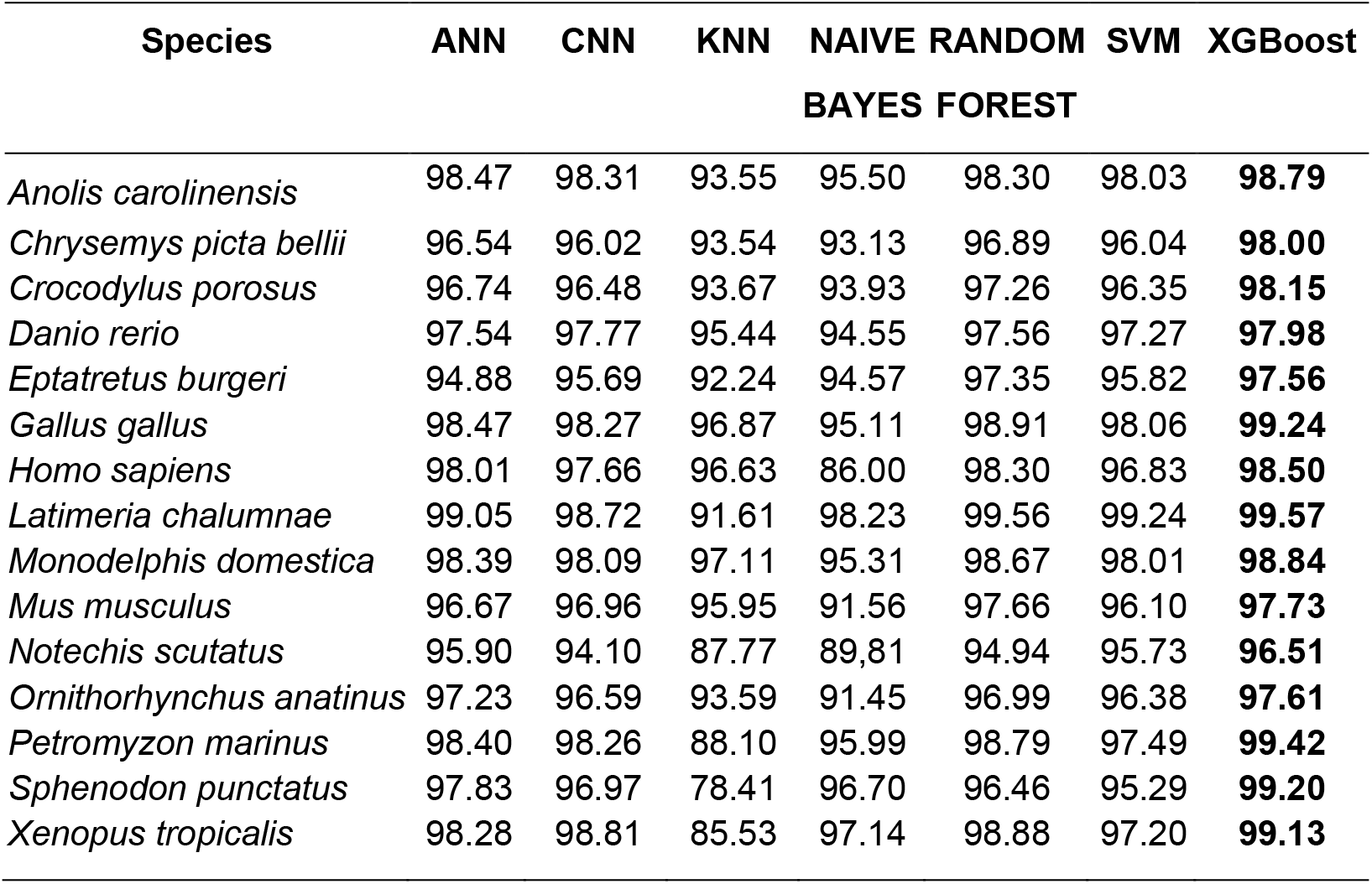
Benchmarking machine learning methods for coding potential prediction based on trinucleotides counts. F1-score for each one of the 15 species in which the algorithms were tested. Other metrics (sensitivity, specificity, precision, accuracy and the confusion matrix) used for the comparison of the algorithm’s performance were made available at the Supplementary File S2.

### Using RNAmining in evolutionary related and unrelated organisms

To demonstrate the generalization of the model built in our tool, we evaluated its performance using the following nine Chordata and non-Chordata organisms that were not used in our training step: *A. thaliana*; *C. elegans*; *C. auratus*; *D. melanogaster*; *G. gorilla gorilla*; *P. textilis*; *R. norvegicus*; *S. cerevisiae; Terrapene carolina triunguis.* In the training set described in the previous topic, we used sequences from representative species from amphibians, birds, mammals, fishes and reptiles. In this new experiment we executed tests using other chordates, but also covering other evolutionary groups such as plants, fungi, insects and nematodes. The F1-score obtained values varying from 86.25 to 98.10. The worst performance was when we used the training set from *L. chalumnae* (Sarcopterygii, Coelacanth) to predict the coding potential of known coding genes and ncRNAs from *D. melanogaster* (Insecta, Diptera). However, the best performance was obtained when we applied the training set from *C. picta bellii* (Sauria, Testudines) in coding and ncRNA sequences from *Terrapene carolina triunguis* (Sauria, Testudines). The F1-score for each organism, together with the respective training set evaluated, can be found in Table 3, meanwhile the confusion matrix and the other metrics can be visualized in Supplementary File S3.

**Table 3.** Evaluation (F1-score) of the models generated by XGBoost, the method implemented in RNAmining, according to evolutionary related and unrelated organisms. Each line comprises the model for each one of the trained species, meanwhile the columns represent the set of 9 evolutionary related and unrelated organisms in which the method was evaluated. Other metrics (sensitivity, specificity, precision, accuracy and the confusion matrix) used for the comparisons were made available at the Supplementary File S3.

| Testing<br>Training | Arabidopsis<br>thaliana | Caenorhabditis<br>elegans | Carassius<br>auratus | Drosophila<br>melanogaster | Gorilla<br>gorilla | Pseudonaja<br>textilis | Rattus<br>norvegicus | Saccharomyces<br>cerevisiae | Terrapene<br>carolina triunguis |
| --- | --- | --- | --- | --- | --- | --- | --- | --- | --- |
| *Anolis carolinensis* | 95.35 | 89.97 | 94.77 | 97.16 | 95.17 | 96.56 | 96.74 | 93.07 | 95.83 |
| *Chrysemys picta bellii* | 97.24 | 97.79 | 95.97 | 98.13 | 97.01 | 97.73 | 97.15 | 96.09 | 98.10 |
| *Crocodylus porosus* | 96.19 | 96.76 | 95.73 | 97.87 | 97.01 | 96.90 | 97.25 | 95.07 | 97.56 |
| *Danio rerio* | 96.64 | 90.50 | 95.29 | 97.96 | 97.24 | 96.89 | 96.42 | 93.96 | 96.62 |
| *Eptatretus burgeri* | 94.90 | 95.57 | 94.80 | 96.73 | 95.34 | 95.43 | 95.76 | 91.49 | 95.51 |
| *Gallus gallus* | 97.60 | 97.89 | 95.76 | 98.02 | 97.93 | 97.79 | 97.59 | 96.48 | 97.69 |
| *Homo sapiens* | 95.71 | 81.25 | 92.19 | 96.44 | 97.73 | 96.24 | 94.60 | 93.57 | 95.65 |
| *Latimeria chalumnae* | 93.71 | 96.78 | 91.63 | 86.25 | 96.30 | 93.39 | 94.37 | 95.47 | 95.63 |
| *Monodelphis domestica* | 97.40 | 97.91 | 95.69 | 98.04 | 97.90 | 97.53 | 97.46 | 93.54 | 97.31 |
| *Mus musculus* | 96.44 | 87.68 | 94.66 | 97.17 | 97.57 | 97.31 | 96.67 | 94.32 | 96.30 |
| *Notechis scutatus* | 97.16 | 97.54 | 95.22 | 97.46 | 97.35 | 97.37 | 96.79 | 94.96 | 97.22 |
| *Ornithorhynchus anatinus* | 97.39 | 97.48 | 95.39 | 87.74 | 97.32 | 97.86 | 97.29 | 94.67 | 97.53 |
| *Petromyzon marinus* | 93.31 | 94.48 | 92.07 | 87.74 | 95.81 | 93.47 | 94.72 | 92.48 | 95.56 |
| *Sphenodon punctatus* | 94.00 | 97.07 | 91.94 | 86.89 | 96.60 | 93.95 | 94.12 | 95.02 | 95.81 |
| *Xenopus tropicalis* | 93.46 | 96.65 | 91.53 | 84.86 | 95.51 | 93.68 | 93.16 | 94.42 | 95.02 |

Even without using any plant in the original training set, we applied the different models to predict the coding potential of known coding and ncRNA sequences from *A. thaliana* (Plantae, Eudicots). The lowest F1-score that RNAmining obtained was 93.31 using a fish model (*Petromyzon marinus,* Agnatha, Petromyzontiformes). The best F1-score was obtained with a marsupial model (*M. domestica,* Marsupialia) that reached 97.40. Thus, this experiment demonstrated the efficiency of the method and the models created even when applied in organisms phylogenetically distant from those used in training.

Finally, in order to show that the results obtained were not by chance, we created 10 datasets of artificial sequences containing the same number, length and nucleotides composition of the coding and ncRNA sequences from the 15 species used in our testing shown in Table 1. The F1-score mean, minimum and maximum values of the 10 datasets from each organism can be visualized in Table 5. The confusion matrix and all the other metrics (accuracy, specificity, sensitivity and precision) can be found in Supplementary File S4. As we can visualize, the F1 measurement mean remained below 38.00 for all artificial sequences created for the tested organisms, with the exception of *P. marinus* (F1-score equals to 64.13), which still had a F1-score below to the values obtained with the other organisms tested for the coding potential prediction (Table 4).

**Table 4.** Evaluation of RNAmining performance according to different sets of artificial sequences from each trained model. F1-score metrics for 10 datasets of artificial sequences randomly generated for each species. The mean, minimum and maximum values are displayed separated by organism. Other metrics (sensitivity, specificity, precision, accuracy and the confusion matrix) used for the comparisons were made available at the Supplementary File S4.

| Species | MEAN | MINIMUM | MAXIMUM |
| --- | --- | --- | --- |
| *Anolis carolinensis* | 1.66 | 0.86 | 2.44 |
| *Chrysemys picta bellii* | 1.08 | 0.70 | 1.40 |
| *Crocodylus porosus* | 0.95 | 0.43 | 1.72 |
| *Danio rerio* | 1.25 | 0.12 | 2.21 |
| *Eptatretus burgeri* | 2.31 | 0.90 | 3.51 |
| *Gallus gallus* | 2.48 | 1.88 | 2.89 |
| *Homo sapiens* | 11.15 | 10.53 | 11.52 |
| *Latimeria chalumnae* | 24.86 | 21.95 | 27.03 |
| *Monodelphis domestica* | 1.34 | 1.00 | 1.18 |
| *Mus musculus* | 6.64 | 5.74 | 7.58 |
| *Notechis scutatus* | 1.80 | 0.58 | 3.99 |
| *Ornithorhynchus anatinus* | 3.62 | 2.67 | 5.04 |
| *Petromyzon marinus* | 64.13 | 62.99 | 65.76 |
| *Sphenodon punctatus* | 37.43 | 31.72 | 41.84 |
| *Xenopus tropicalis* | 23.26 | 17.65 | 28.21 |

### Comparing RNAmining performance with publicly available tools

Next, we compared RNAmining performance with other four tools commonly used for nucleotides coding potential prediction: CPAT, CPC2, RNAcon and Transdecoder. We used as input all coding and ncRNA sequences from the testing dataset used in the 15 species listed in Table 1. According to the F1-score metric, RNAmining outperformed all the tools in all organisms with the exception of CPAT for *L. chalumnae*, in which both tools presented an equal F1-score of 99.57. The comparative performance of all tools can be observed in Table 5. The detailed results regarding their accuracy, sensitivity, specificity, precision, F1-score and the confusion matrix can be found in Supplementary File S2. Finally, we used the t-student test to compare the results from RNAmining and the other tools, revealing that our software presented significantly better results in performing coding potential predictions based on known coding genes and ncRNAs. The p-values obtained in these comparisons were: 0.0026 (*vs* CPAT); 1.57e-05 (*vs* CPC2); 2.69e-05 (*vs* RNAcon); and 2.89e-05 (*vs* Transdecoder).

**Table 5.** Benchmarking results from RNAmining and the other tools already described in the literature according to organisms from different evolutionary branches. F1-score metric for CPAT, CPC2, RNAcon, Transdecorder and RNAmining, based on the predictions using models provided by each tool or generated according to their instructions. The bold numbers are the best values regarding F1-score metric. The results for other metrics were made available at the Supplementary File S2.

| Species | CPAT | CPC2 | RNAcon | Transdecoder | RNAmMining |
| --- | --- | --- | --- | --- | --- |
| *Anolis carolinensis* | 94.55 | 86.87 | 83.03 | 88.26 | **98.79** |
| *Chrysemys picta bellii* | 92.56 | 89.01 | 82.36 | 84.80 | **98.00** |
| *Crocodylus porosus* | 94.07 | 92.48 | 84.32 | 87.63 | **98.15** |
| *Danio rerio* | 94.64 | 87.17 | 80.97 | 87.74 | **97.98** |
| *Eptatretus burgeri* | 95.59 | 78.82 | 75.84 | 76.26 | **97.56** |
| *Gallus gallus* | 96.95 | 90.69 | 75.81 | 83.50 | **99.24** |
| *Homo sapiens* | 95.20 | 75.85 | 71.73 | 76.02 | **98.50** |
| *Latimeria chalumnae* | **99.57** | 91.60 | 97.45 | 98.86 | **99.57** |
| *Monodelphis domestica* | 96.24 | 91.44 | 80.90 | 85.22 | **98.84** |
| *Mus musculus* | 95.48 | 81.40 | 76.78 | 80.80 | **97.73** |
| *Notechis scutatus* | 85.19 | 86.29 | 84.83 | 83.44 | **96.51** |
| *Ornithorhynchus anatinus* | 87.47 | 72.04 | 84.73 | 84.63 | **97.61** |
| *Petromyzon marinus* | 96.59 | 75.14 | 95.11 | 96.68 | **99.42** |
| *Sphenodon punctatus* | 97.61 | 91.91 | 97.86 | 95.24 | **99.20** |
| *Xenopus tropicalis* | 99.07 | 97.92 | 98.70 | 97.77 | **99.13** |

### RNAmining stand-alone and web server tool

RNAmining tool was made available in both stand-alone and web server versions. The tools only require the nucleotide sequences of the RNAs in which the user intends to perform the coding potential prediction in FASTA format, together with the species name in a standardized format related to the model to be used. Besides our tool presented good results even when using phylogenetically distant organismos, we recommend to always use the most closely-related species to the one the user wants to perform the predictions. Furthermore, RNAmining documentation presents all the guidelines on how to generate a model for a particular set of sequences and organisms of interest. The web interface of RNAmining tool was developed to allow users to quickly perform the coding potential prediction without the need of installing any specific program and using only a generic internet browser. The only requirement for running the tool is a FASTA file containing the nucleotide sequences and the organism model that the user wants to use, which can be selected in a drop-down menu containing all 15 organisms used in the training step (Figure 2A). There is no limit of the number of sequences, but the web server supports files up to 20Mb. For files bigger than that, we recommend to use the stand-alone RNAmining tool. RNAmining will automatically classify the FASTA sequences used as input and identify which of them are coding or non-coding RNAs. Finally, as a result it offers a table with the sequences’ IDs and its classification as coding or non-coding, which can also be downloaded in tabular format, together with two separate FASTA files containing both the coding and non-coding sequences separately (Figure 2B).

**Figure 2.**
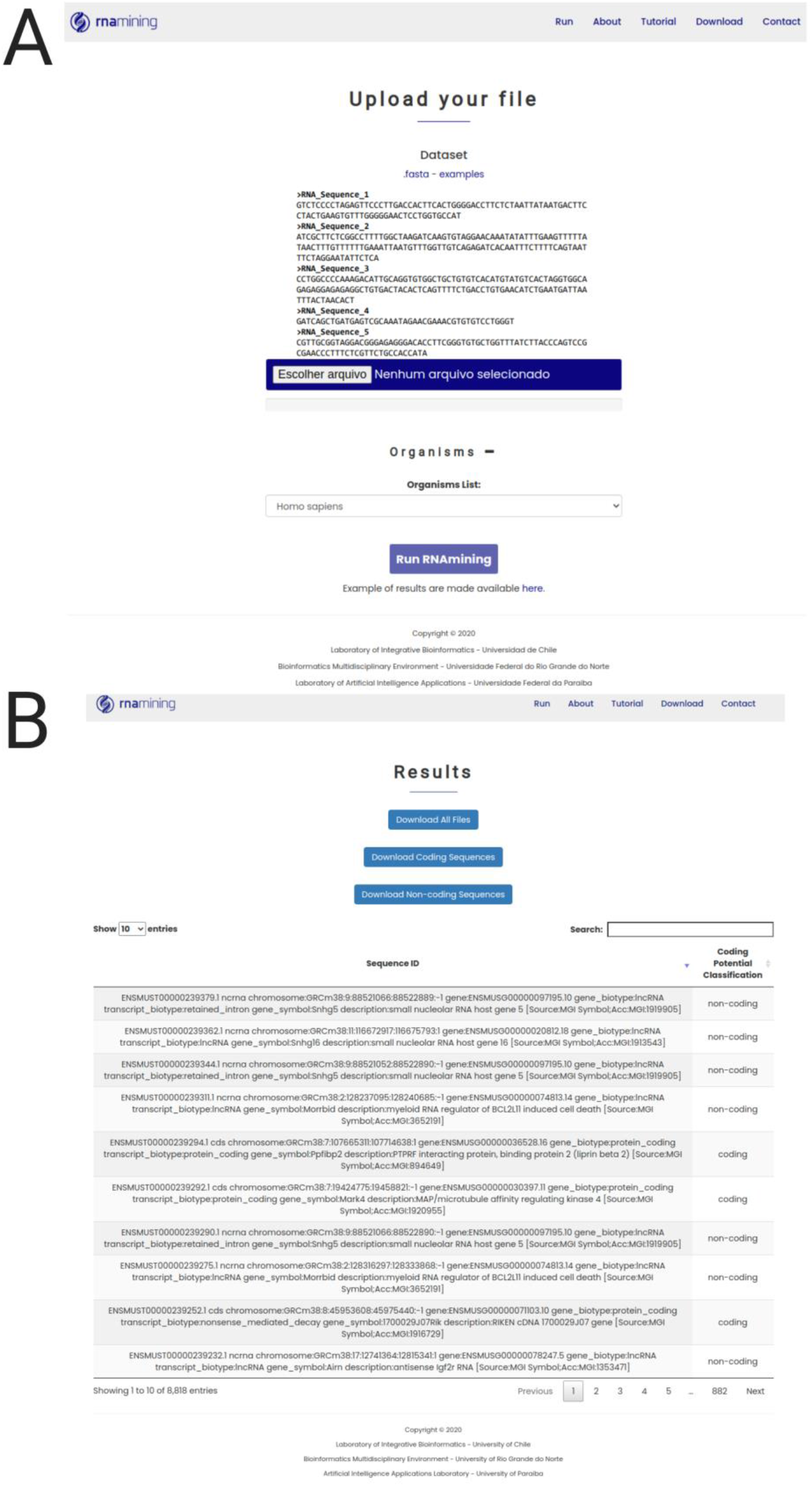
RNAmining web server overview. **A. Job launcher screen (Run tab)**. The user only needs to upload the nucleotide sequences in FASTA format and select the model to be used based on the evolutionary close related species. **B. Results web page screen**. General report containing the list of coding and non-coding sequences in a dynamic table, in which the user can search for a particular sequence or filter only those coding or non-coding RNAs by using a free text form that will filter the results in the table dynamically. The user can download the complete table in tabular format and two FASTA files containinng the set of coding and non-coding RNAs separately.

## DISCUSSION

The coding potential prediction of nucleotides is a key step in the definition of the repertoire of noncoding RNAs in a genome or transcriptome project, especially when dealing with non-model organisms. Sometimes, predictive tools for the computational characterization of RNA molecules, in analyses like the prediction of specific RNA families (19) or the estimation of a network of RNA-RNA (20) or protein-RNA interactions (21), have their performance affected according to the training organism, increasing the number of false positives when applied in evolutionarily distant species. In this work, we evaluated the performances of seven different supervised machine learning algorithms, using eukaryotic species from a variety of evolutionary clades, revealing their potential to be used in the development of novel and improved computational tools for the coding potential prediction of RNA sequences. Artificial intelligence has been widely used in computational biology (22,23), but its application to characterize ncRNAs has been limited.

In this benchmarking, we opted to analyze the tri-nucleotides counting as the main feature to be evaluated for the coding potential prediction, followed by a normalization considering the sequences length (*i.e.* each tri-nucleotide count was divided by the total size of the given sequence). Panwar *et al.* (10) used nucleotides counting successfully for this purpose. They considered 40,905 non-coding RNAs from Rfam release 10.0 database and 62,473 coding RNA sequences from Human RefSeq database, divided into 50% of training and 50% of test (*i.e*. the training and test sets were composed of 20,453 non-coding and 31,237 coding sequences). They used the counts of mono-, di-, tri-, tetra- and penta-nucleotides and a combination of all counts using the SVM method, and showed that using tri-nucleotides counts is enough to predict the coding potential of ncRNAs with better accuracies. Our comparisons of the machine learning algorithms revealed XGBoost as the algorithm with better performance, presenting efficiency in predicting the coding potential of RNA sequences even when using the models of distantly related organisms. This latter shows the usefulness of this approach for performing coding predictions in non-model organisms.

We implemented XGBoost in RNAmining, a stand-alone and web server tool flexible to be used in genome or transcriptome projects focused in both model and non-model eukaryotic organisms. Our tool outperformed similar approaches, such as CPAT (11), CPC2 (13), RNAcon (10) and Transdecoder (12). Both versions of the software are easy to use, with the web version providing a simple report and FASTA format files that can be used in downstream analysis. It provides 15 models generated from eukaryotic from different evolutionary clades. Other models can be generated by the user using the stand-alone version, which can be used with simple command line operations. These features facilitate its usage for experienced users and, especially, for those without any programming experience, which can easily perform large-scale predictions of the coding potential of nucleotide sequences in both genome or transcriptome initiatives.

## DATA AVAILABILITY

Ensembl is an open access genome browser for vertebrate genomes in the Ensembl website (https://www.ensembl.org/index.html).

RNAmining is a tool for coding potential prediction which is freely available at (https://rnamining.integrativebioinformatics.me/download).

## SUPPLEMENTARY DATA

Supplementary Data are available at NAR online.

## ACKNOWLEDGEMENT

The authors would like to thank Dr. Savio Torres de Farias for the helpful discussions during the preparation of this manuscript.

